# Phylogenetic parallelograms: visual comparison of discordant phylogenetic trees

**DOI:** 10.64898/2026.09.18.752558

**Authors:** Daniel H. Huson, Banu Cetinkaya, Louxin Zhang

## Abstract

Phylogenetic trees inferred from different genomic regions, or from organellar and nuclear genomes, frequently disagree owing to incomplete lineage sorting, hybridization, introgression or horizontal gene transfer. Such discordance is informative, yet the standard tool for comparing trees visually, the tanglegram, is restricted to two trees and is known to misrepresent their similarity. Here we introduce phylogenetic parallelograms, drawings in which any number of rooted trees are embedded into a common scaffold, a rooted phylogenetic network that displays all of them, and are drawn in parallel so that shared branches align while conflicting branches visibly diverge. On synthetic tree pairs, the complexity of a parallelogram tracks the topological distance between the trees, whereas the complexity of an optimized tanglegram does not. We apply the approach to genome-wide introgression in the *Anopheles gambiae* species complex and to organellar discordance in cats, grasses and Fagaceae. PhyloParallelograms is an open-source interactive application.

## Introduction

The evolutionary history of a group of species is usually summarized as a single phylogenetic tree, but trees estimated from different parts of the genome frequently disagree. Incomplete lineage sorting and, more interestingly, processes such as hybridization, introgression and horizontal gene transfer, can cause the history of an individual locus to depart from that of the species carrying it^1–4^. Phylogenomic studies routinely estimate hundreds or thousands of gene trees, and the pattern of their disagreement has itself become an object of study. Gene tree discordance is used to detect introgression across a genome ^5–7^, to identify chromosomal regions with distinct histories, such as inversion regions or sex chromosomes ^7;8^, and to document discordance between organellar and nuclear genomes in animals and plants ^9–13^. In each of these settings, a researcher needs to see not only that trees disagree, but where they disagree, which clades are involved, and which alternative topologies recur across loci.

A number of approaches are used to summarize discordance. Concordance factors and related quantities ^14–16^ count, for each branch of a reference tree, how many gene trees support or contradict it; they quantify conflict but do not display the alternative topologies. Embeddings of tree space^17^ show the distances between trees while hiding the trees themselves. DensiTree ^18^ superimposes many trees on common leaf positions and is effective for posterior samples of similar trees, but for a moderate number of distinct topologies the overlay becomes difficult to interpret. Consensus and split networks^19;20^ display the incompatible splits found in a collection of trees, at the cost of losing the identity of the individual trees. Finally, rooted phylogenetic networks inferred from gene trees ^21–24^ represent explicit hypotheses of reticulate evolution. However, they do not show how the input trees are related to each other.

The standard tool for comparing two trees directly is the tanglegram^25–30^, in which the trees are drawn facing each other and corresponding leaves are connected by lines. Tanglegrams are intuitive, widely used and available in many phylogenetic viewers^29;31;32^. However, arranging two trees so as to minimize the number of line crossings is computationally hard^33;34^, and, more fundamentally, the number of crossings is a poor indicator of topological difference. De Vienne ^35^ showed by simulation that trees differing in many branches can usually be drawn without any crossing at all, so that a tanglegram can give a misleading impression of congruence. Tanglegrams are also inherently pairwise and do not lend themselves to the comparison of more than two trees.

Here we introduce phylogenetic parallelograms, a visualization in which any number of rooted phylogenetic trees are drawn together in a single coordinate system, with corresponding branches drawn in parallel wherever the trees agree. The construction rests on a simple idea. Rather than aligning the trees to each other directly, we first compute a scaffold, a rooted phylogenetic network that displays every input tree, using the PhyloFusion algorithm ^24^. Each tree is then traced through the scaffold and drawn as a slightly offset copy of the scaffold edges that it uses. Where the trees agree, their branches form bundles of parallel lines; where a tree places a clade differently, its branch leaves the bundle and re-enters the drawing elsewhere, and every such departure corresponds to a reticulation of the scaffold. The number of reticulations, the hybridization number of the scaffold, thereby provides a natural measure of the complexity of a parallelogram. We show on synthetic tree pairs that this measure tracks the topological distance between two trees, whereas the complexity of an optimized tanglegram does not. To illustrate our new approach, we apply parallelograms to genome-wide introgression in the *Anopheles gambiae* species complex^7^ and to discordance between organellar and nuclear trees in the living cats ^11^, danthonioid grasses ^13^ and the oak family Fagaceae ^12^, on trees with six to 129 taxa. The method is implemented in PhyloParallelograms, an open-source interactive application.

## Results

### Phylogenetic parallelograms

Figure 1 introduces phylogenetic parallelograms using two trees for six species of the *Anopheles gambiae* complex, taken from Fontaine et al. ^7^: the species tree, which is supported by the distal part of the X chromosome, and the tree obtained from the whole genome, which is dominated by autosomal loci that have experienced extensive introgression. In the tanglegram (Fig. 1a) the leaves of the two trees line up perfectly and no connecting line crosses another, so that one may easily conclude that the trees are nearly identical. In fact, they differ substantially. In the whole-genome tree, *An. arabiensis* is sister to the *An. coluzzii*–*An. gambiae* pair rather than to *An. quadriannulatus*, and *An. melas* and *An. merus* form a clade rather than successive lineages below the other species. The corresponding phylogenetic parallelogram (Fig. 1b) makes these differences immediately visible. The two trees are drawn in different colours in a shared coordinate system. Branches on which the trees agree, such as those leading to the *An. coluzzii*–*An. gambiae* pair and to *An. melas* and *An. merus*, run in parallel, whereas each disagreement appears as a branch that leaves the shared structure and attaches elsewhere.

**Figure 1:**
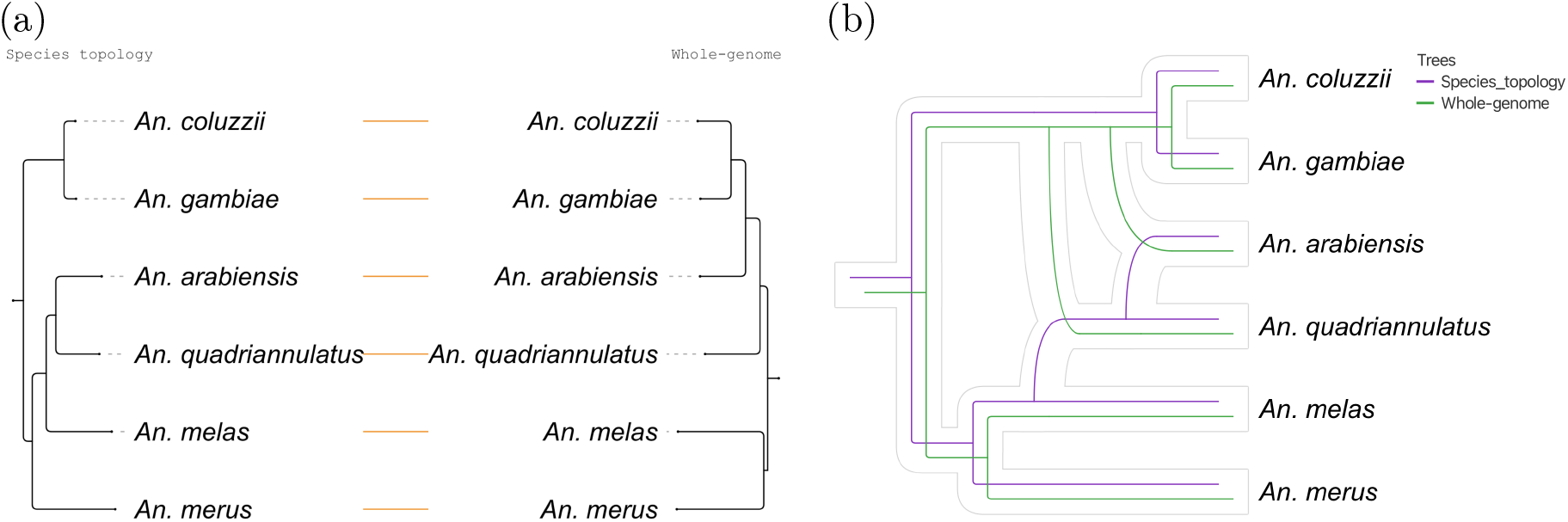
Tanglegram versus phylogenetic parallelogram. (a) Tanglegram comparing the species tree and the whole-genome tree for six species of the *Anopheles gambiae* complex, following Fontaine et al. ^7^. Corresponding leaves are connected by lines; the drawing has no crossings. (b) Phylogenetic parallelogram computed from the same two trees. Both trees are drawn in a shared coordinate system, in different colours. Branches on which the trees agree run in parallel; each branch that leaves the shared structure and attaches elsewhere marks a clade that one tree places differently. The underlying network is indicated here as an outline.

Reading a parallelogram requires only a few conventions. All trees share one ordering of the taxa and one root. A bundle of parallel lines represents a branch present in all trees of the corresponding colours, so that the width of a bundle shows how many trees share a branch. A line that leaves a bundle, typically drawn as a curve, and joins the drawing at another position marks a clade that the corresponding tree places differently, and the position at which the line joins shows where. Every such departure corresponds to a reticulation of the underlying scaffold, and the number of reticulations, the hybridization number of the scaffold, summarizes the amount of conflict in the drawing. A taxon that is absent from a tree simply has no line of that tree’s colour. Unlike a tanglegram, the drawing does not depend on the pairing of trees, and unlike a rooted phylogenetic network, every input tree remains individually visible. The name is a play on words: a parallelogram is a collection of cladograms or phylograms drawn in parallel.

### Computing a phylogenetic parallelogram

A phylogenetic parallelogram is computed in four steps (Fig. 2; Methods). First, a scaffold network *N* that displays all input trees is computed using PhyloFusion ^24^. PhyloFusion encodes each tree as a set of lineage taxon strings, aligns these strings across trees by computing shortest common supersequences, and reads a rooted network off the alignment. The algorithm is a heuristic for minimizing the hybridization number of *N*, a computationally hard problem^36^. It accepts multifurcating trees and trees with missing taxa, and it produces tree-child networks ^37^, in which every internal node has at least one child that is not a reticulate node. Second, for every node and every reticulate edge of *N* we record the set of input trees whose embedding passes through it. These tree-membership annotations arise naturally in the alignment step, in which occurrences of the same taxon in different trees are merged (Fig. 2, subscripts), and they are completed by a single post-order traversal of *N*. Third, branch lengths of the input trees, when present, are transferred to the scaffold by a non-negative least-squares fit, so that phylograms as well as cladograms can be drawn. Fourth, a layout of *N* is computed using the algorithms of PhyloSketch^38^, which place the reticulate edges of a rooted network so as to minimize their displacement, and each input tree is drawn as a copy of the scaffold edges that carry its index, shifted by a small tree-specific offset (Fig. 2, bottom right). Trees that were not used to compute the scaffold can subsequently be added to the drawing by an exhaustive search over the reticulate edges of *N* (Methods). This allows a user to compute a scaffold from a subset of trees and then display any further tree that the scaffold happens to contain.

**Figure 2:**
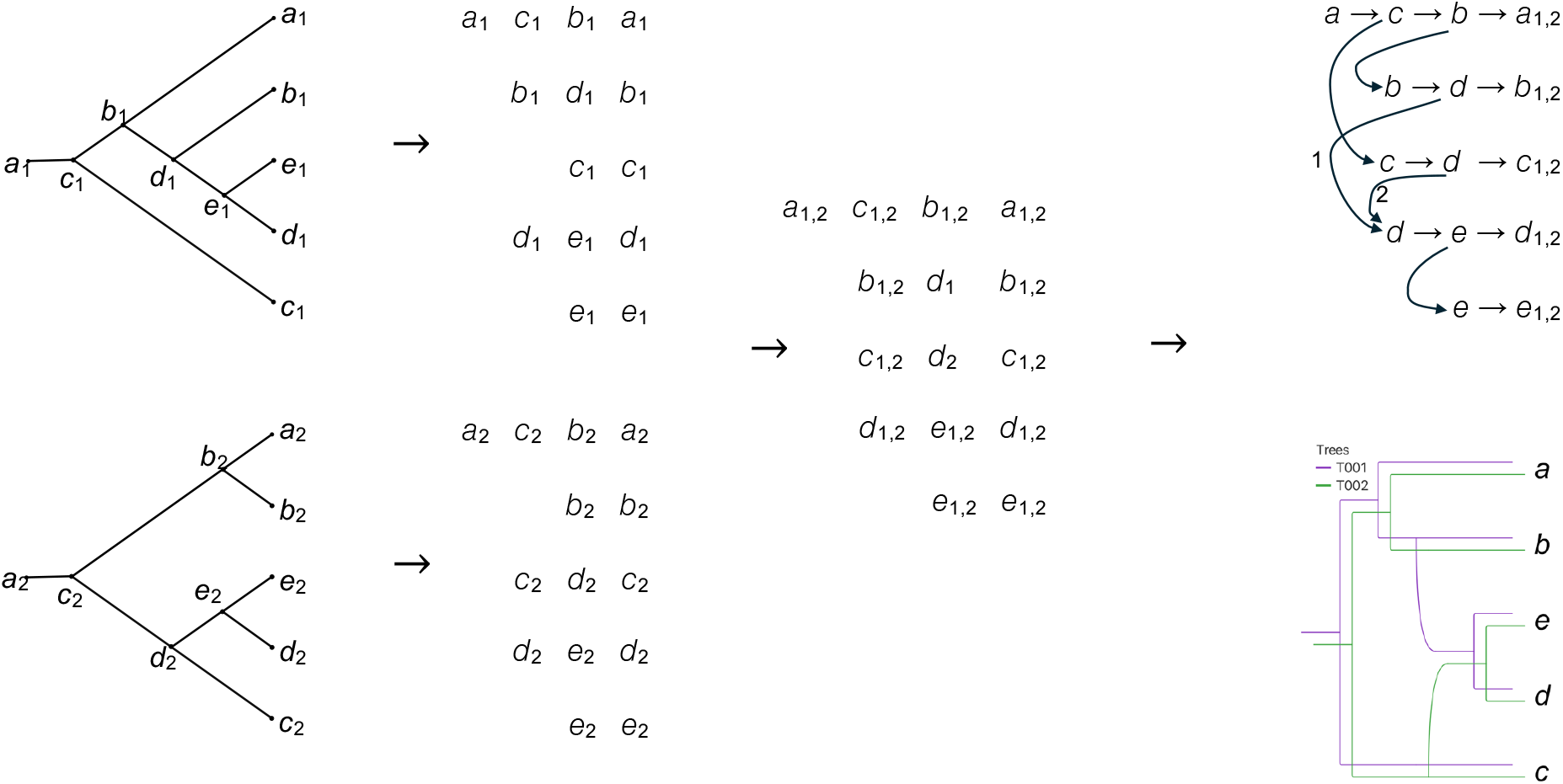
Computing a scaffold by encoding, aligning and decoding. From left to right: two input trees *T*_1_ (top) and *T*_2_ (bottom) on *X* = {*a*, …, *e* }, labelled under the ordering *a < b < c < d < e* so that every taxon occurs once on an internal node and once on a leaf; the lineage taxon string (LTS) encoding of each tree, one string per taxon; for each taxon, the shortest common supersequence (SCS) of its two strings, which differ only for *b* and *c*; and the rooted network obtained by decoding the SCS encoding, which displays both trees and has a reticulate node wherever a taxon is reached from two or more lineages, here *d*. Subscripts give the tree of origin: each occurrence in an LTS string carries the index of its tree and, in the SCS encoding, merged occurrences carry both indices, whereas the occurrence of *d* in the string of *b* comes from *T*_1_ alone and that of *d* in the string of *c* from *T*_2_ alone. These two occurrences induce the two reticulate edges, labelled 1 and 2 in the network, in which every leaf carries both indices because both trees contain all five taxa. Here, only the leaves are labelled, to avoid clutter. Bottom right: the phylogenetic parallelogram of *T*_1_ and *T*_2_ drawn on this network.

The scaffold is a computational device rather than an evolutionary hypothesis. Its reticulations localize the differences among the trees, but they are not claims about hybridization or gene flow. Nevertheless, because *N* displays every input tree with a heuristically minimal number of reticulations, the scaffold is closely related to the hybridization networks that have been studied for two decades ^21;39;40^, and a parallelogram can be viewed as such a network annotated with the exact path of every input tree.

### Parallelogram complexity reflects topological distance

To examine how faithfully the two visualizations reflect the actual difference between two trees, we used the synthetic tree pairs of de Vienne ^35^, who generated pairs of trees on 20 taxa at Robinson–Foulds (RF) distances ^41^ of 2, 4, …, 34 and showed that the minimum number of crossings in a tanglegram is only weakly related to the RF distance. Figure 3 shows four of these pairs, with RF distances of 2, 12, 26 and 32, drawn both as displacement-optimized tanglegrams ^30^ and as phylogenetic parallelograms. The first three tanglegrams can be drawn without any crossing, even though the trees at RF distance 26 share only a minority of their clades; only the most divergent pair requires crossings, and only four of them. The parallelograms, in contrast, grow progressively more complex: the pair at RF distance 2 differs by a single reticulation, whereas the pairs at RF distances 12, 26 and 32 require 4, 8 and 10 reticulations, respectively.

**Figure 3:**
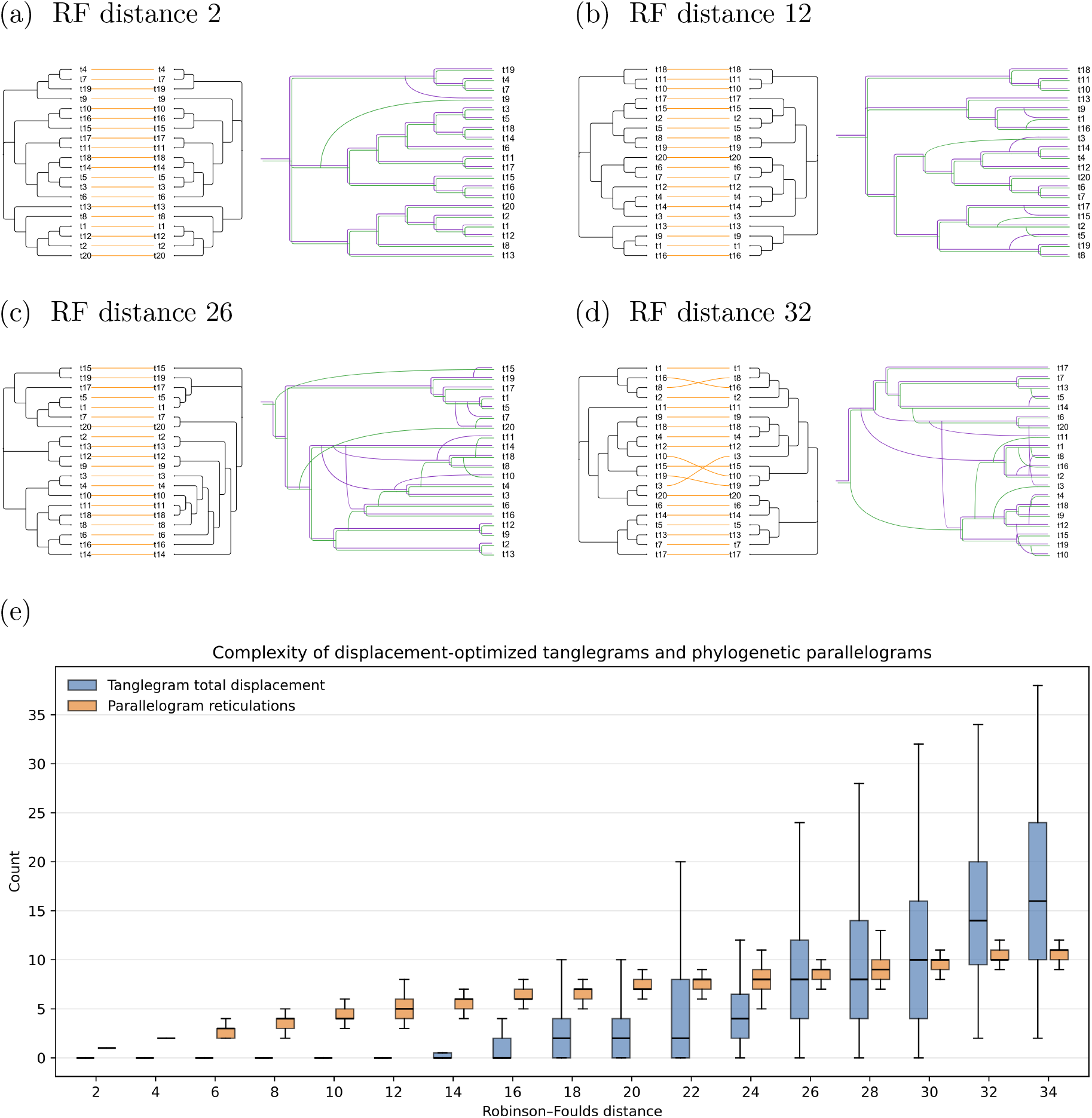
Tanglegrams and phylogenetic parallelograms for synthetic tree pairs. (a)–(d) Displacement-optimized tanglegrams ^30^ (left) and phylogenetic parallelograms (right) for four pairs of synthetic trees on 20 taxa from the simulation study of de Vienne ^35^, with increasing Robinson–Foulds distances. The first three tanglegrams have no crossings, whereas the corresponding parallelograms reveal progressively larger amounts of disagreement. (e) For each Robinson–Foulds distance of 2, 4, …, 34, box plots summarize, over 100 pairs of synthetic trees on 20 taxa ^35^, the minimum total displacement of a displacement-optimized tanglegram ^30^ (blue) and the hybridization number of the scaffold network underlying the phylogenetic parallelogram (orange). Boxes show the median and interquartile range; whiskers extend to minimum and maximum. Source data are provided as a Source Data file.

We then quantified this behaviour over the complete data set of 100 tree pairs for each of the 17 RF distances (Fig. 3e). For each pair we computed the minimum total displacement of a tanglegram, that is, the smallest achievable sum of vertical offsets between corresponding leaves^30^, and the hybridization number of the scaffold underlying the parallelogram. For RF distances up to 12, every one of the 600 optimized tanglegrams has zero displacement, and the median displacement remains at or below four up to an RF distance of 24. The displacement then rises steeply, but with very large variation among pairs of equal RF distance, spanning values from 2 to 38 at RF distance 34. The hybridization number, by contrast, increases steadily from 1 at RF distance 2 to a median of 11 at RF distance 34, and its interquartile range rarely exceeds two. The complexity of a phylogenetic parallelogram is thus a far more consistent visual proxy for topological difference than the complexity of a tanglegram, and it remains informative in exactly the regime of moderate RF distances in which tanglegrams fail. Hybridization number and RF distance measure different things and need not coincide: a single rooted subtree-prune-and-regraft move creates one reticulation but can change many clades.

### Recurring alternative histories across the *Anopheles* genome

Fontaine et al. ^7^ showed that the species of the *Anopheles gambiae* complex have exchanged genes so extensively that most of the genome supports a tree different from the species tree, which is recovered only from the distal part of the X chromosome, and that particular chromosomal regions, notably the 2La and 3La inversions, carry yet other histories. To illustrate the use of parallelograms for genome-wide comparisons, we selected 15 loci of 3–7 kb from the published whole-genome alignment (Methods): five from the distal X chromosome, one from the pericentromeric X chromosome, five from the autosomal arms 2R and 3R, and two each from the 2La and 3La inversions (Table 1). We inferred a maximum-likelihood tree for each locus and rooted the trees using *An. christyi* as outgroup.

**Table 1:**
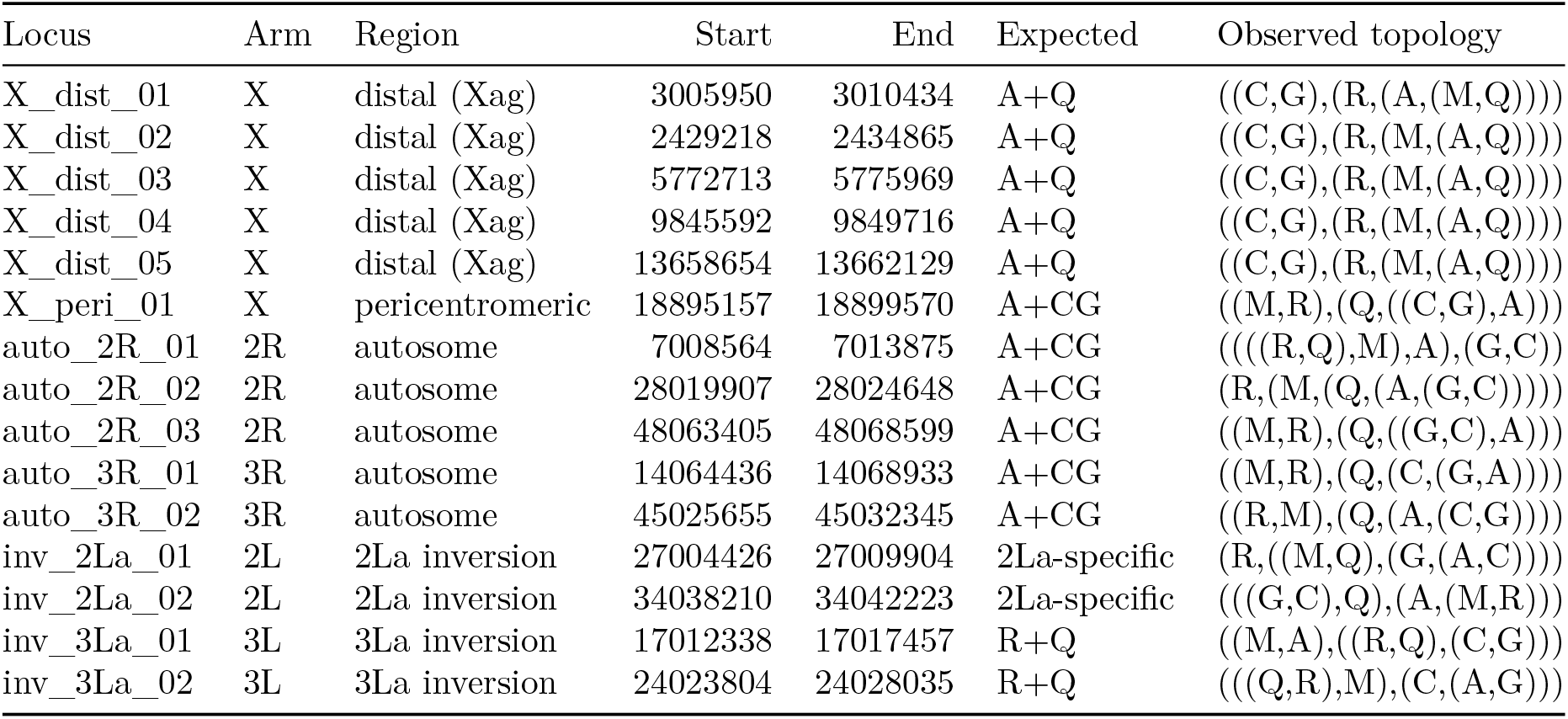
Loci used for the *Anopheles* gene trees. Coordinates refer to the *An. gambiae* PEST assembly that anchors the whole-genome alignment of Fontaine et al. ^7^. The expected topology follows Fontaine et al.: A+Q, *An. arabiensis* sister to *An. quadriannulatus* (species tree); A+CG, *An. arabiensis* sister to *An. coluzzii* and *An. gambiae*; R+Q, *An. merus* with *An. quadriannulatus*. The last column gives the rooted ingroup topology of the maximum-likelihood tree (A, *An. arabiensis*; C, *An. coluzzii*; G, *An. gambiae*; M, *An. melas*; Q, *An. quadriannulatus*; R, *An. merus*).

The parallelogram of all 15 trees (Fig. 4) is drawn with the five X-distal trees highlighted. Four of them have exactly the topology of the species tree in Fig. 1, and the fifth differs from it only in placing *An. quadriannulatus* as sister to *An. melas*, a grouping with 47% bootstrap support. These trees form the thick bundle that runs through the drawing and against which the remaining trees can be read. None of the other ten trees has the species-tree topology. The pericentromeric X locus and two of the autosomal loci reproduce the whole-genome tree of Fig. 1 exactly, and two further autosomal loci share its defining feature, a clade uniting *An. arabiensis* with *An. coluzzii* and *An. gambiae*, while differing in detail: one places *An. melas* and *An. merus* as successive rather than sister lineages, and the other places *An. arabiensis* as sister to *An. gambiae* alone. In the parallelogram, all five of these trees converge on the reticulate edge that carries *An. arabiensis* to the *An. coluzzii*–*An. gambiae* lineage. Three trees, both loci from the 3La inversion and one 2R locus, group *An. merus* with *An. quadriannulatus* (85–94% bootstrap support), consistent with the introgression between these species that Fontaine et al. reported for this region, and their branches share the reticulate edge that attaches *An. merus* to *An. quadriannulatus*. The two loci from the 2La inversion, finally, support histories not seen elsewhere in the sample: one groups *An. arabiensis* with *An. coluzzii* to the exclusion of *An. gambiae* with 100% bootstrap support and places *An. melas* with *An. quadriannulatus*, whereas the other, poorly supported, tree attaches *An. arabiensis* to *An. melas* and *An. merus*.

**Figure 4:**
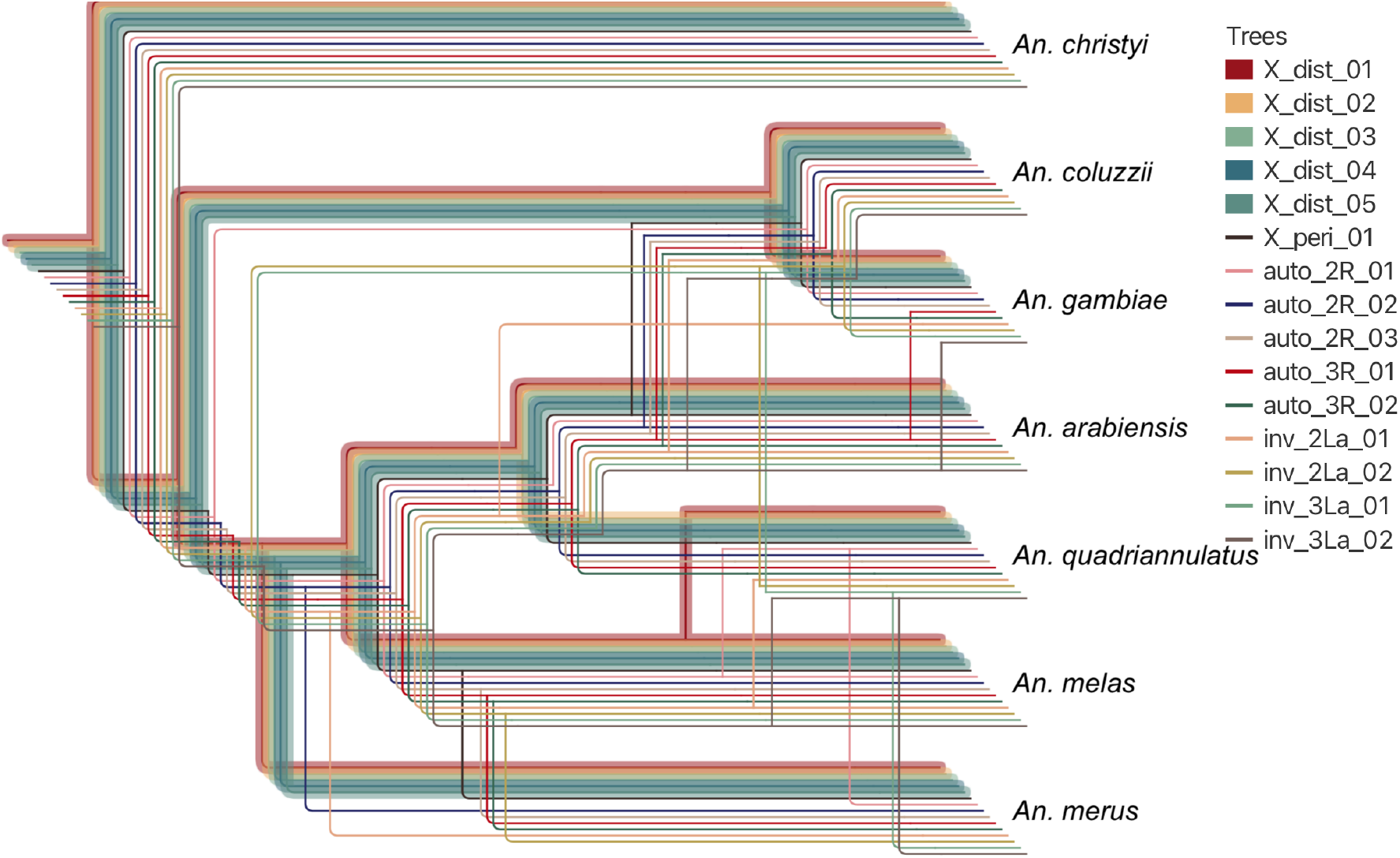
Phylogenetic parallelogram of 15 gene trees from different regions of the *Anopheles gambiae* complex genome. Trees were inferred from loci in the distal (X_dist) and pericentromeric (X_peri) X chromosome, the autosomal arms 2R and 3R (auto), and the 2La and 3La inversions (inv), with *An. christyi* as outgroup. The five X-distal trees, which are expected to reflect the species tree, are highlighted in bold. Reticulate edges are drawn in transfer-edge style: at each reticulate node, the incoming edge used by the majority of trees is layed out as a tree edge and the remaining incoming edges as transfer edges.

While summary methods may struggle with such data, the parallelogram displays the conflict, and does so in a way that reveals which loci share which alternative history. Two features of the software are useful for such collections. Because Fig. 4 shows 15 trees, the reticulate edges are drawn in the transfer-edge style of PhyloSketch^38^: at each reticulate node, the incoming edge used by the majority of trees is layed out as a tree edge and the others as transfer edges, which keeps the dominant signal legible. Furthermore, the set of displayed trees can be changed interactively without recomputing the layout, so that, for example, the three trees that group *An. merus* with *An. quadriannulatus* can be shown on their own against the X-distal bundle.

### Organellar discordance at three scales

Discordance between organellar and nuclear phylogenies is among the most frequently reported forms of conflict in both animals ^9^ and plants ^10^, and pairs of large trees are the situation in which tanglegrams are most often used. We compared organellar and nuclear trees from three studies of increasing size and difficulty.

Li et al. ^11^ inferred the phylogeny of cats from genome-wide nuclear data and from complete mitochondrial genomes and attributed the differences between the two trees to ancient hybridization. We transcribed both trees from their Fig. 1A (Methods). The nuclear tree has 42 tips, including three samples of the Asian leopard cat and the Sunda clouded leopard, which are absent from the mitogenome tree of 39 tips, so that the trees share 38 taxa and five taxa occur in only one of them. The tanglegram (Fig. 5a) has a single crossing, between the domestic cat and the African wild cat, again suggesting broad agreement. The parallelogram (Fig. 5b) shows that the trees differ at *h* = 9 positions. For example, the two trees disagree on the placement of the Pallas cat, and of the caracal lineage.

**Figure 5:**
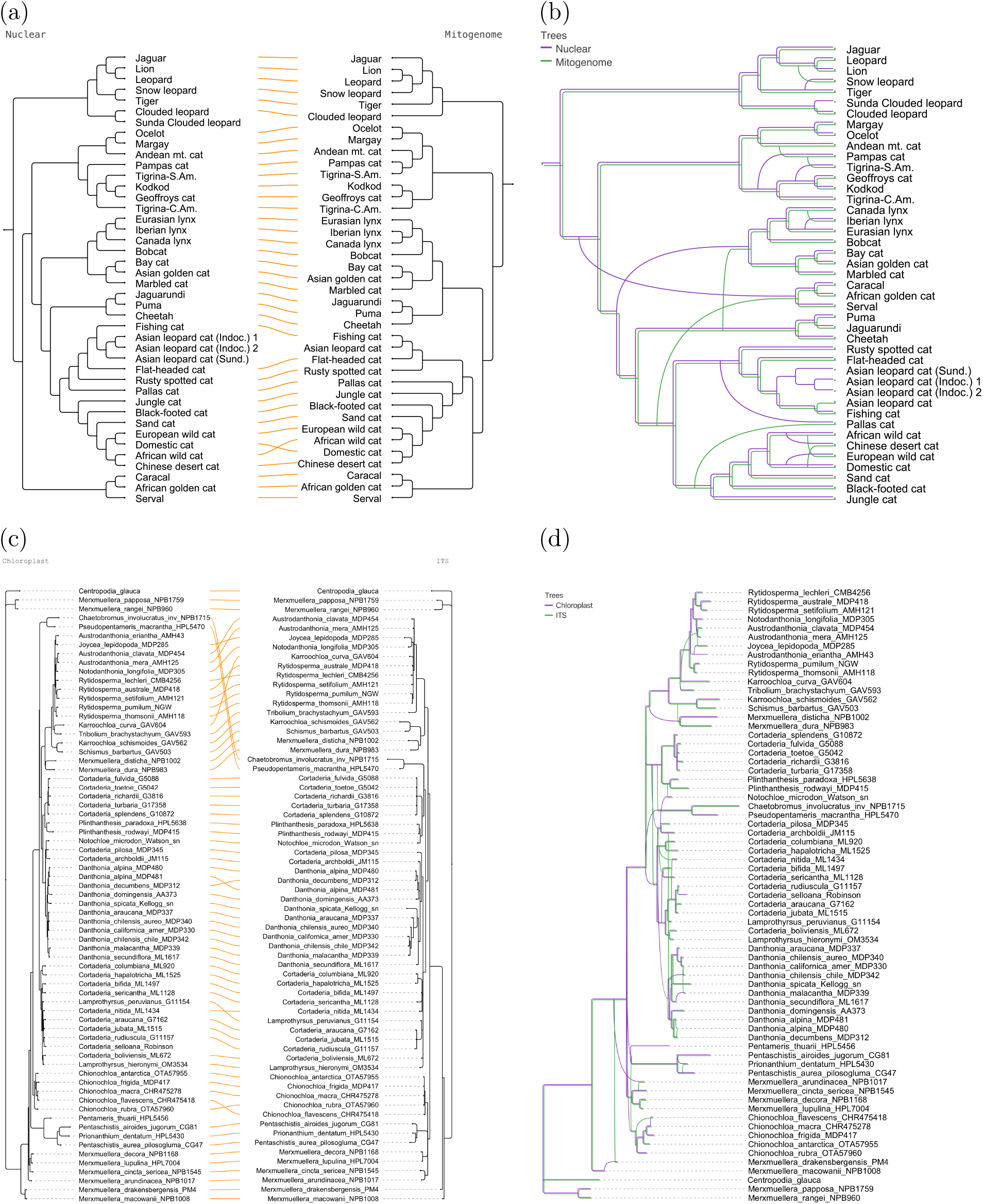
Organellar discordance in cats and in danthonioid grasses. (a) Tanglegram comparing the nuclear and mitogenome trees of cats, transcribed from Li et al. ^11^. (b) Phylogenetic parallelogram computed from the same two trees; the four taxa that occur only in the nuclear tree and the Asian leopard cat, which occurs only in the mitogenome tree, appear as branches of a single colour. (c) Tanglegram comparing chloroplast and ITS trees for 69 taxa of Danthonioideae, adapted from Pirie et al. ^13^. (d) Phylogenetic parallelogram computed from the same two trees. Panels (a) and (b) use cladograms, which show topology only; panels (c) and (d) use phylograms, in which branch lengths are drawn to scale.

The chloroplast and ITS phylogenies of the grass subfamily Danthonioideae by Pirie et al. ^13^comprise 69 taxa (Fig. 5c,d). Here the tanglegram has several crossings. The parallelogram shows that the two markers agree on the backbone of the subfamily and on most genera, and many of the disagreements are localized to relationships within *Danthonia* and within the *Rytidosperma* group.

Finally, Zhou et al. ^12^ inferred nuclear and plastid phylogenies for 129 representatives of Fagaceae and documented extensive cytonuclear discordance, which they attributed to introgression. Here the tanglegram (Fig. 6a) is so heavily tangled that it is difficult to tell which parts of the trees agree. The corresponding parallelogram (Fig. 6b) has *h* = 49 and several congruent regions are readily discernible. This example shows that parallelograms remain readable for trees with more than a hundred taxa and dozens of conflicts.

**Figure 6:**
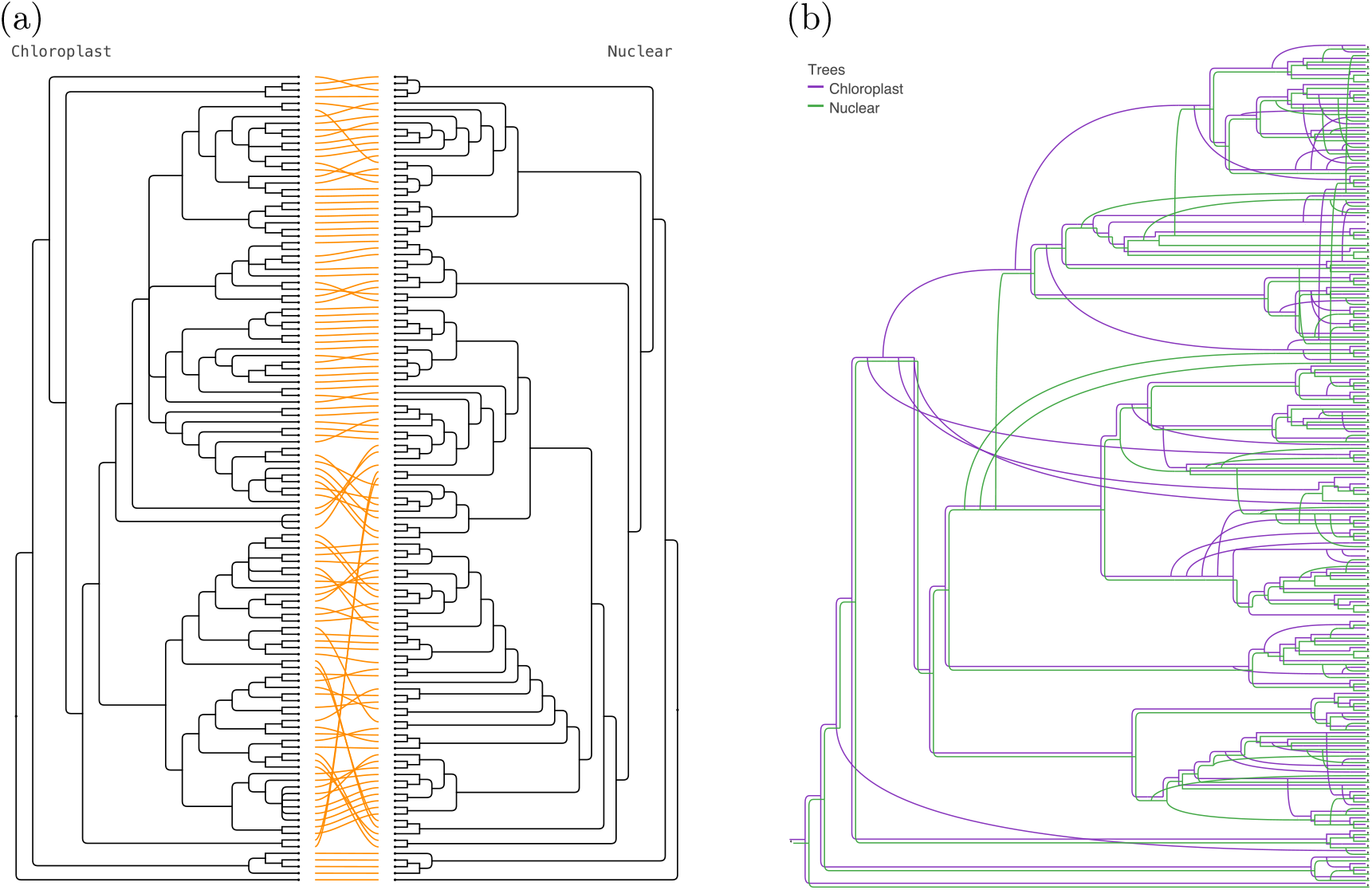
Cytonuclear discordance in Fagaceae. (a) Tanglegram comparing the plastid and nuclear phylogenies of 129 Fagaceae taxa, following Zhou et al. ^12^. (b) Phylogenetic parallelogram computed from the same two trees. Shared structure forms parallel bundles, and the numerous localized conflicts appear as branches that leave and re-enter the shared structure. Taxon labels are omitted for legibility; the labelled trees are available in the data repository.

### Interactive exploration

PhyloParallelograms is a desktop application for macOS, Windows and Linux (Fig. 7). Trees are loaded from Newick or Nexus ^42^ files and listed in a table with two check boxes per tree, one selecting the trees used to compute the scaffold and one selecting the trees that are displayed. Separating these two choices is useful in practice. A scaffold computed from a few well-supported trees provides a stable frame of reference into which further trees can be embedded one at a time, and the display can be restricted to any subset of trees in order to examine a particular conflict. Branches with support below a user-defined threshold can be contracted before the scaffold is computed, which removes reticulations that would otherwise be caused by poorly supported branches; the resulting multifurcations are handled by PhyloFusion, which can also mutually refine the input trees so that a multifurcation in one tree does not conflict with a resolved branch in another ^24^.

**Figure 7:**
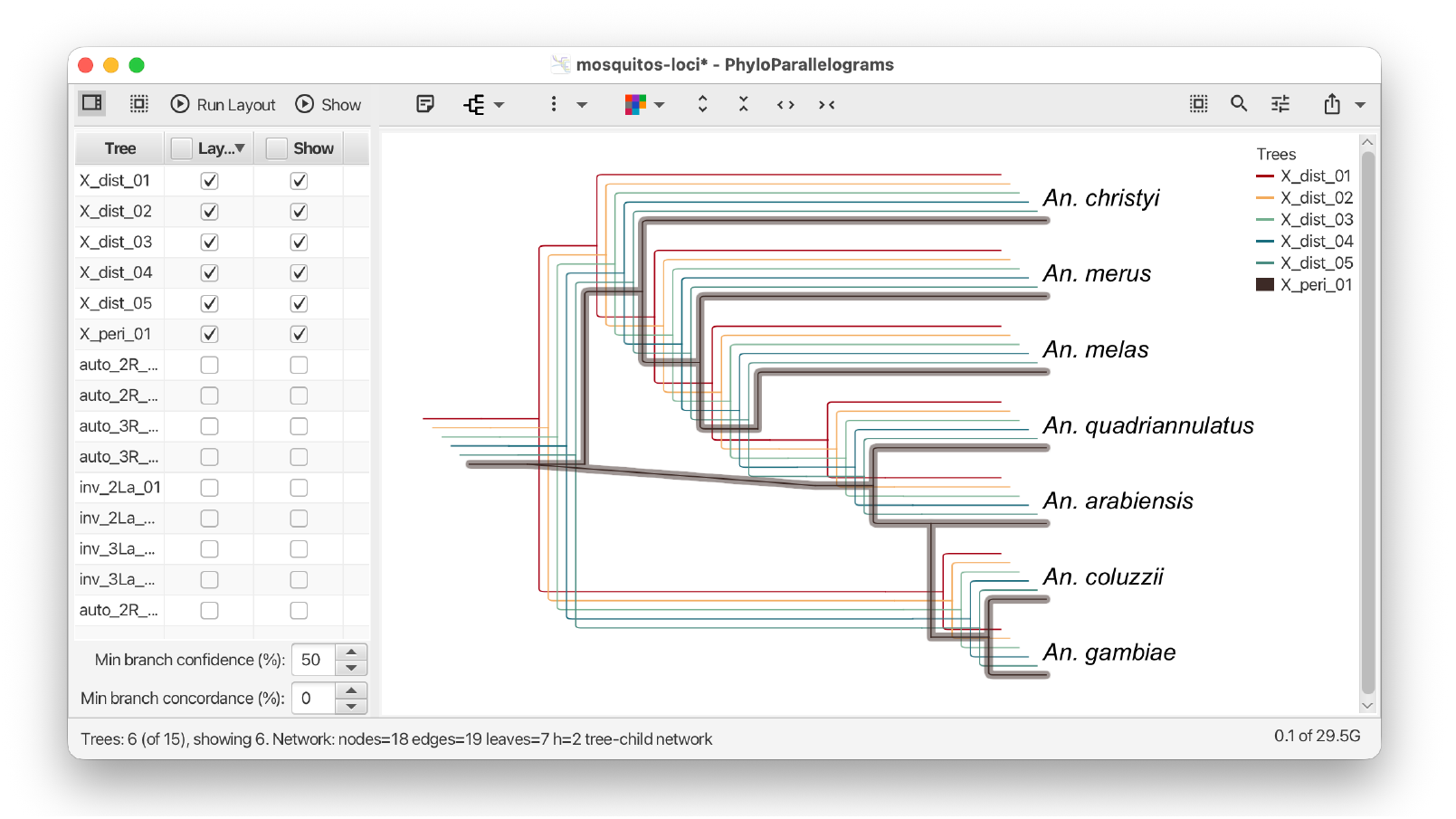
Interactive exploration in PhyloParallelograms. The 15 gene trees of Fig. 4 are loaded. In the tree table (left), only the five X-distal trees and the pericentromeric X tree are selected for computing the scaffold (Layout) and for display (Show). The minimum branch confidence is set to 50%, so that branches with lower bootstrap support are contracted. In the resulting parallelogram (right), the five X-distal trees form a single bundle, from which the pericentromeric tree (shown in bold) clearly departs.

Figure 7 illustrates this on the gene trees of Fig. 4. All 15 trees are loaded, but only the five X-distal trees and the pericentromeric X tree are selected for the scaffold and for display, and the minimum branch confidence is set to 50%. This contracts the three branches with lower bootstrap support among the six trees: the branch (47%) that placed *An. quadriannulatus* with *An. melas* in one X-distal tree, and two branches (41% and 20%) that arranged *An. melas* and *An. merus* in another. The five X-distal trees then agree and form a single bundle, and only the pericentromeric tree departs from it, by attaching *An. arabiensis* to the *An. coluzzii*–*An. gambiae* lineage and by uniting *An. melas* with *An. merus*.

Trees can be rerooted by outgroup or midpoint, taxa can be removed, and the parallelogram can be drawn as a rectangular, circular or radial cladogram or phylogram, with reticulate edges drawn straight or curved, or in the transfer-edge style used in Fig. 4. Trees are coloured according to a selectable colour scheme, individual trees can be renamed, recoloured and highlighted, and drawings are exported in PNG, SVG, PDF or LATEX format, together with the scaffold in extended Newick format. An analysis is saved in a text-based document format that records the input trees, the settings, the scaffold and its tree-membership annotations. Several example data sets, including the mosquito and Danthonioideae data used here, are distributed with the software.

### Performance

Computing a parallelogram is dominated by the construction of the scaffold. PhyloFusion has been evaluated in detail elsewhere^24^ and handles tens of trees on hundreds of taxa in seconds to minutes. For all data sets shown here, the complete computation took only seconds on a laptop computer (Table 2), and rendering is linear in the total size of the tree embeddings. The practical limit of the approach is not the number of trees or taxa but the amount of conflict among the trees: a scaffold with many reticulations yields a crowded drawing regardless of how it is laid out, and in such cases the support filter and the selection of subsets of trees are the appropriate tools.

**Table 2:** Data sets used in this study. For each data set, the number of taxa and trees, the hybridization number *h*(*N*) of the scaffold network and the wall-clock time for computing the parallelogram on a laptop computer (MacBook Pro, Apple M4) are given.

| Data set | Source | Taxa | Trees | $h(N)$ | Time (s) |
| --- | --- | --- | --- | --- | --- |
| *Anopheles*, species and whole-genome trees | ref. 7 | 6 | 2 | 2 | 0.1 |
| *Anopheles*, gene trees from 15 loci | ref. 7 | 7 | 15 | 7 | 0.1 |
| Cats, nuclear and mitogenome trees | ref. 11 | 43 | 2 | 9 | 0.1 |
| Danthonioideae, chloroplast and ITS trees | ref. 13 | 69 | 2 | 21 | 4 |
| Fagaceae, plastid and nuclear trees | ref. 12 | 129 | 2 | 49 | 8 |
| Synthetic tree pairs (17 RF distances × 100 pairs) | ref. 35 | 20 | 2 | 1–12 | 0–1 |

## Discussion

Phylogenetic parallelograms address a gap between the two families of tools that biologists currently use to look at discordant trees. Tanglegrams compare two trees but can hide their differences, whereas networks and summary statistics condense many trees into one object but discard the trees themselves. A parallelogram keeps every tree visible and legible, aligns the trees where they agree, and localizes their disagreements to specific clades, for any number of trees that share a scaffold of manageable complexity. The drawings can be read without training: a bundle is agreement, a curve is conflict, and the colour says which tree.

The synthetic comparison makes precise what is otherwise a matter of impression. The number of reticulations required to accommodate two trees increases smoothly with their RF distance and varies little among trees of equal distance, whereas the displacement of an optimal tanglegram is zero for a wide range of substantially different trees and then varies wildly. The reason is structural. A tanglegram can absorb any rearrangement that preserves the order of the leaves, however many clades it destroys, whereas a parallelogram must account for every clade that is not shared.

Several caveats apply. The scaffold is a heuristic solution to a hard optimization problem, and different scaffolds displaying the same trees, with the same or a slightly different number of reticulations, may lead to different but equally valid parallelograms; where a reticulation is placed reflects a choice among alternatives that the software exposes as an option (Methods). The reticulations of the scaffold mark where trees differ, not why, and they should not be read as hybridization events without further analysis. A rooted tree is required for every input, which for gene trees means that an outgroup or a rooting method has to be chosen. Finally, the hybridization number, like the RF distance, treats all conflicts alike, whereas a biologist may care more about some clades than about others; the interactive selection of trees and taxa is intended to compensate for this.

Several extensions suggest themselves. In a parallelogram, every reticulate edge carries a weight, namely the number of trees that use it, which defines a weighted version of the displacement-minimization problem solved by PhyloSketch; solving this weighted problem should improve the layout of parallelograms with many trees. We are also exploring a filter that contracts branches of the input trees that appear to reflect incomplete lineage sorting rather than reticulation. For collections of hundreds of gene trees, clustering trees by the reticulate edges that they use, or computing a scaffold from a small number of representative trees and then embedding the remainder, are natural strategies that the current implementation already supports in part. We also envisage parallelograms of rooted networks, in which the network from one analysis is embedded in the scaffold of another. Phylogenetic parallelograms are available in the open-source application PhyloParallelograms, and we anticipate that they will find use wherever gene trees, marker trees or trees produced by different methods need to be compared.

## Methods

### Rooted phylogenetic trees and networks

Let *X* be a set of taxa. A rooted phylogenetic tree *T* on *X*^*′*^ ⊆ *X* is a rooted tree whose leaves are labelled bijectively by *X*^*′*^; internal nodes may have more than two children (multifurcations), which arise, for example, when branches with low support are contracted. Every edge *e* of *T* defines a cluster, the set of taxa below *e*, and *T* is determined by its set of clusters. A rooted phylogenetic network *N* on *X* is a directed acyclic graph with a single root, in which all leaves are labelled bijectively by *X*, and in which every node other than the root either has in-degree one (a tree node) or in-degree greater than one (a reticulate node). Edges entering a reticulate node are called reticulate edges. The hybridization number of *N* is *h*(*N*) = Σ_*v*_(indeg(*v*) −1), where the sum runs over all reticulate nodes *v. N* is tree-child if every internal node has at least one child that is not a reticulate node ^37^. *N* displays a tree *T* on *X*^*′*^ ⊆ *X* if *T* can be obtained from *N* by deleting all but one incoming edge of every reticulate node, deleting all nodes that do not lie on a path from the root to a leaf labelled by an element of *X*^*′*^, and suppressing all nodes with in-degree and out-degree one. We use display in the strict sense that the resulting tree is *T* itself, including its multifurcations, rather than a resolution of *T*. The subgraph of *N* traversed in this way is the embedding of *T* in *N*. Computing a network with minimum hybridization number that displays a given set of trees is NP-hard ^36^.

### Computation of the scaffold

Given input trees *T*_1_, …, *T*_*k*_, the scaffold is a rooted phylogenetic network *N* that displays all of them, computed with PhyloFusion^24^, an extension of the ALTS algorithm^40^ to multifurcating trees and unequal taxon sets. Briefly, based on a fixed ordering of the taxa, PhyloFusion assigns to every node of every tree a taxon label such that each taxon appears exactly twice in each tree, once on an internal node and once on a leaf (Fig. 2, left). Each tree is thereby encoded as a set of lineage taxon strings, one per taxon (Fig. 2, second column). The strings of all trees are aligned by computing shortest common supersequences (Fig. 2, third column), using heuristics described in ref. 24, and the resulting alignment induces a tree-child network that displays all input trees (Fig. 2, top right). The network obtained in this way depends on the taxon ordering, over which the algorithm searches heuristically in order to minimize *h*(*N*); for the two trees of Fig. 2, the ordering *a < b < c < d < e* yields one reticulation, whereas *a < b < e < d < c* yields two.

The labelling and the strings have a simple interpretation (Fig. 8). Build a tree one taxon at a time in the given order: start with the root, labelled *x*_1_, and the leaf *x*_1_ attached to it, and attach each further leaf *x*_*i*_ by subdividing an edge of the tree built so far, giving the new node the label *x*_*i*_. Every internal node arises in exactly one such step, so every taxon labels exactly one internal node and one leaf; equivalently, an internal node is labelled by the larger of the smallest taxa below its two children. The lineage taxon string of *x*_*i*_, read from the node labelled *x*_*i*_ down to the leaf *x*_*i*_, lists the taxa that were subsequently attached to the lineage of *x*_*i*_, in order from the root. A tree is thus represented as a set of lineages, one per taxon, each recording which later taxa branch off from it and in which order, and two trees are compared lineage by lineage. If both trees attach the same taxa in the same order to the lineage of *x*, their strings for *x* coincide; otherwise the shortest common supersequence of the two strings is the shortest lineage that accommodates both, and every occurrence of a taxon beyond its first, summed over all strings, costs one reticulation. In Fig. 2, the lineages of *a, d* and *e* are identical in both trees, whereas *T*_1_ attaches *d*, and with it *e*, to the lineage of *b* and *T*_2_ attaches it to the lineage of *c*; the supersequences retain both attachments, and decoding therefore reaches the node labelled *d* from two lineages.

**Figure 8:**
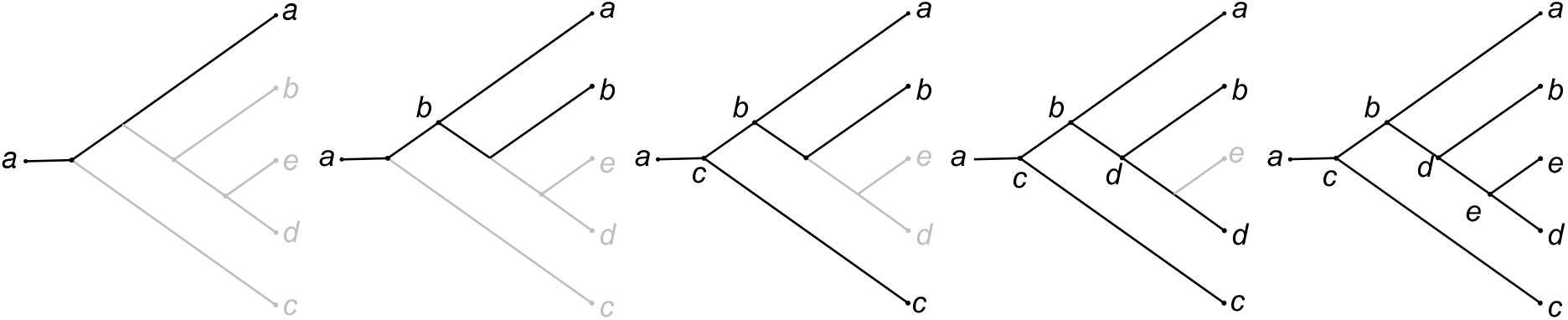
Building the node labelling one taxon at a time. The tree *T*_1_ of Fig. 2 is built up under the ordering *a < b < c < d < e*; the part still to be added is shown in grey. The tree starts as the root, labelled *a*, with the leaf *a* attached to it. Each further taxon is attached by subdividing an edge of the tree built so far, and the new node receives that taxon’s label. Every taxon therefore labels exactly one internal node and one leaf, and the lineage taxon string of a taxon *x*, read from the node labelled *x* to the leaf *x*, lists the taxa that were subsequently attached to the lineage of *x*.

Several preprocessing steps are optional. Edges of the input trees with support below a user-defined threshold are contracted. Taxa that are never separated by any input tree can be grouped, which reduces the problem size without changing the result. The input trees can be mutually refined, so that a multifurcation in one tree that is resolved in a compatible way by another tree is resolved accordingly, which removes reticulations that would only reflect different degrees of resolution. A missing-taxa heuristic uses the trees that contain a taxon to construct a compatible lineage for it in trees in which it is missing. When several networks with the same number of reticulations are available, the user can choose which subnetwork to place below a reticulate node: the default avoids reticulate edges that connect a node to one of its own descendants, and alternatives place the subnetwork with the smallest or the largest number of taxa below the reticulate node.

### Tree-membership annotations

For every node *v* and every reticulate edge *e* of *N*, we record sets *τ* (*v*), *τ* (*e*) ⊆ {1, …, *k*} of the indices of the input trees whose embeddings pass through *v* and *e*, respectively. The annotations are introduced during the alignment step of PhyloFusion. Each occurrence of a taxon in a lineage string (Fig. 2, second column) carries the identifier of the tree from which it originates. When the shortest common supersequence is computed (Fig. 2, third column), aligned occurrences of the same taxon are merged and their tree identifiers are combined by set union. Each position of the supersequence becomes a node of *N*, and its set of identifiers is the initial annotation *τ* (*v*). Reticulate edges are induced by individual taxon occurrences within aligned positions and inherit the identifiers of these occurrences as their initial annotation *τ* (*e*). The subscripts in Fig. 2 show these identifiers: all positions of the SCS encoding are contributed by both trees except two. The occurrence of *d* in the string of *b* stems from *T*_1_ only, and the occurrence of *d* in the string of *c* from *T*_2_ only. These two positions give rise to the two reticulate edges entering the node labelled *d*, which represents the clade {*d, e*}, and the edges are used by *T*_1_ and *T*_2_, respectively. In the scaffold, after completion of the annotations as described next, every node is annotated with {1, 2}, whereas each of the two reticulate edges is used by exactly one tree, which is the annotation drawn in Fig. 2; accordingly, the parallelogram there shows the clade {*d, e*} attached below *b* for *T*_1_ and below *c* for *T*_2_.

The initial annotations cover only the nodes and reticulate edges that are created directly from the aligned positions. The complete embedding of every input tree is then obtained in a single post-order traversal of *N*. A leaf is annotated with the identifiers of all trees that contain the corresponding taxon. For an internal node *v, τ* (*v*) is the union of the contributions of its outgoing edges, where a tree edge (*v, w*) contributes *τ* (*w*) and a reticulate edge (*v, w*) contributes *τ* (*v, w*). The resulting annotations associate every node and reticulate edge with exactly those input trees whose embeddings pass through it, and the embedding of *T*_*i*_ consists of all nodes *v* with *i ∈ τ* (*v*), all tree edges (*v, w*) with *i ∈ τ* (*v*) *∩ τ* (*w*), and all reticulate edges *e* with *i ∈ τ* (*e*).

For multifurcating trees, annotations are maintained for individual taxon occurrences rather than for entire elements of the lineage strings, because a multifurcation contributes several taxa to the same element while alignment is performed separately for each occurrence; different taxa from the same multifurcation may therefore become associated with different subsets of trees, and different reticulate edges entering the same node may carry different annotations. When a taxon that is missing from a tree is supplied by the missing-taxa heuristic, the newly introduced occurrence retains the identifiers of the tree from which it was copied, so that the tree lacking the taxon is not attributed a branch that it does not contain. The annotations are written into the extended Newick description of the scaffold (as node and edge attributes named TT), so that a scaffold together with its embedded trees can be saved and exchanged.

### Embedding additional trees

A tree *T* that was not used to compute the scaffold can be embedded in *N* if *N* happens to display it. This is decided by exhaustive search. Let *C*(*T*) be the set of clusters of *T*. For every combination of choices of one incoming edge per reticulate node, extract the tree *T* ^*′*^ displayed by *N* under that combination and let *C*(*T* ^*′*^) be the set of its clusters. If the two sets *C*(*T*) and *C*(*T* ^*′*^) are compatible, then record *T* in the annotations *τ* in the same way as for the original input trees.

Note that if *T* has unresolved nodes, then there might be several displayed trees with compatible clusters, in which case *T* will be represented by an embedded network rather than a tree. The number of combinations is the product of the in-degrees of the reticulate nodes, so this procedure is practical for scaffolds with a moderate number of reticulations, which is precisely the regime in which parallelograms are useful. The same search can optionally be applied to the input trees themselves, to show all embeddings in the case of multifurcations.

### Branch lengths

The annotations determine, for each input tree *T*_*i*_, the subset of *N* traversed by its embedding, from which a displayed copy of *T*_*i*_ is extracted. Descendant clusters are computed for the input tree and for this copy, and matching clusters identify, for every edge *e* of *T*_*i*_, the path *P* (*e*) of scaffold edges that represents it. Let *w*(*e*) be the length of *e* in *T*_*i*_ and let *x*_*f*_ be the unknown length of scaffold edge *f*. Scaffold edge lengths are estimated such that Σ_*f*∈*P*(*e*)_ *x*_*f*_ approximates *w*(*e*) for all edges *e* of all input trees. By default, the non-negative least-squares estimate is used, which minimizes the sum of squared residuals subject to *x*_*f*_ ≥ 0. Alternatively, the sum of absolute residuals can be minimized by linear programming, optionally with all reticulate edges constrained to length zero, or each tree-edge length can be distributed evenly along its path and each scaffold edge assigned the mean of the contributions it receives. The optimization problems are solved using the ojAlgo library. Scaffold edge lengths are used when a phylogram is drawn; cladograms do not require them.

### Layout

Coordinates are first assigned to the scaffold *N* using the layout algorithms of PhyloSketch^38^. Leaves are placed in the order in which they occur in the scaffold, tree edges are drawn as in a rectangular, circular or radial cladogram or phylogram, and the reticulate edges are placed so as to minimize their total displacement, the sum of the distances between the endpoints of reticulate edges, which is the criterion used by PhyloSketch for drawing rooted networks. Each input tree *T*_*i*_ is then rendered in the same coordinate system by copying every scaffold edge in the embedding of *T*_*i*_ and translating the copy by a tree-specific offset. The offsets (*x*_*i*_, *y*_*i*_) are distributed evenly along the diagonal from (−*w/*2,−*w/*2) to (*w/*2, *w/*2), where *w* is a user-defined spread parameter (*w* = 10 for all figures in this article except Fig. 4, in which a larger value was used to separate the 15 trees).

As a result, branches shared by several trees are drawn as bundles of parallel lines, whereas branches on which the trees disagree diverge. Reticulate edges can be drawn as straight segments or as curves. In the transfer-edge style used in Fig. 4, the incoming edges of every reticulate node are compared by the number of trees using them; if the most frequently used edge is used by at least a user-defined fraction of these trees (50% by default), it is drawn as a tree edge, and the remaining incoming edges are drawn as horizontal edges in the manner of horizontal gene transfers, which improves legibility when many trees traverse the same reticulation. Trees are assigned colours from a selectable colour scheme, and a legend lists the displayed trees.

The rendering step is linear in the total size of the embeddings, that is, in the sum over all displayed trees of the number of scaffold edges that they use. The computationally demanding steps are the construction of the scaffold and its layout; both are heuristic solutions to hard optimization problems, and both are analysed in the publications describing PhyloFusion ^24^and PhyloSketch^38^, respectively. In a parallelogram, every reticulate edge *e* naturally carries the weight |*τ*(*e*)|, which gives rise to a weighted displacement-minimization problem; the current implementation uses the unweighted PhyloSketch layout, and the weighted problem is left for future work.

### Synthetic comparison with tanglegrams

We used the synthetic tree pairs generated and provided by de Vienne ^35^, consisting of 100 pairs of rooted trees on 20 taxa for each Robinson–Foulds distance 2, 4, …, 34. For each pair, a displacement-optimized tanglegram was computed with the algorithm of ref. 30, as implemented in SplitsTree ^20^, and the minimum total displacement was recorded; the displacement of a tanglegram is the sum, over all taxa, of the vertical distance between the two occurrences of the taxon, so that a displacement of zero corresponds to a tanglegram in which all connecting lines are horizontal. In parallel, the scaffold network for the pair was computed with PhyloFusion using default settings, and its hybridization number was recorded. The Robinson–Foulds distances were taken from the data set. This batch computation was performed using a shell script and a SplitsTree workflow, see Code availability.

### *Anopheles* data

Whole-genome alignments of the *An. gambiae* complex in MAF format were obtained from the Dryad repository associated with Fontaine et al.^7^ (doi:10.5061/dryad.f4114). Candidate loci were selected from long alignment blocks that contain all species of interest. To identify suitable regions, we scanned the chromosome-specific MAF files, detected alignment blocks with complete species coverage, and selected windows of several kilobases that fall entirely within a single such block, which avoids artefacts caused by stitching together fragmented or partially covered alignments. In this way we selected 15 loci of 3.3–6.7 kb: five in the distal region of the X chromosome, one in the pericentromeric region of the X chromosome, three on 2R, two on 3R, two within the 2La inversion on 2L and two within the 3La inversion on 3L (Table 1). For each locus, the aligned sequences were extracted with the `MafIndex` functionality of Biopython ^43^, species identifiers were mapped to species names, and one alignment in FASTA format was written per locus; the reference sequence *An. gambiae* PEST, which anchors the alignment, and the second outgroup *An. epiroticus* were excluded, giving alignments of the six species of the complex and *An. christyi* as outgroup. Coverage, alignment length and gap fractions were computed for each alignment as quality-control statistics. Maximum-likelihood trees were inferred separately for each locus with IQ-TREE 3^44^ (version 3.1.2) using the command `iqtree3 -redo -S data/alignments -B 1000 -T AUTO -st DNA`. All trees were rooted on *An. christyi*. The species tree and the whole-genome tree shown in Fig. 1 were transcribed from Fig. 1B of Fontaine et al. ^7^. A step-by-step description of the preparation of these data, including the scripts used for locus selection and extraction, is available at https://github.com/husonlab/trees-to-networks-tutorial.

### Organellar and nuclear data sets

The nuclear and mitogenome trees of the living cats were transcribed from Fig. 1A of Li et al. ^11^ with the image-capture function of PhyloSketch ^38^, after removing all elements of the figure other than the tree edges and the leaf labels and converting the coloured and dashed edges to solid black lines. The nuclear tree comprises 42 tips, including two Indochinese and one Sundaic sample of the Asian leopard cat and the Sunda clouded leopard, and the mitogenome tree 39 tips, with a single Asian leopard cat; the five taxa that occur in only one tree were retained, and the scaffold was computed with the missing-taxa procedure of PhyloFusion. Both trees were rooted as in the original figure.

The chloroplast and ITS trees of Danthonioideae were transcribed in the same way from Figure 1 of Pirie et al. ^13^ and rooted as in that publication. Transcription was necessary because neither study provides the trees as files: Li et al. deposited the underlying sequence and genotype data only, and the data link given by Pirie et al. is apparently no longer accessible.

The nuclear and plastid phylogenies of Fagaceae were recomputed in MrBayes v3.2.6 ^45^ from the data sets provided by Zhou et al. ^12^, using the partitioning schemes and substitution models of the supplied NEXUS files, *Betula pendula* as outgroup, and the run settings reported in their paper (10 million generations, sampling every 100 generations, first 25% discarded as burn-in), which differ from the settings embedded in the supplied files. The trees used here are the all-compatible consensus trees with posterior probabilities. Because the two data sets use different taxon labels, the labels were matched by hand using the sample information provided by Zhou et al., and labels that could not be matched were left unchanged.

### Tanglegrams

All tanglegrams shown in this article were computed with the displacement-optimized tanglegram algorithm ^30^ as implemented in SplitsTree ^20^.

### Software implementation

PhyloParallelograms is written in Java using JavaFX and builds on the jloda3 and SplitsTree6 libraries ^20^, which provide the PhyloFusion algorithm and the tree and network data structures, and the layout code used in PhyloSketch ^38^. The application reads and writes Newick and Nexus ^42^ files, exports drawings in PNG, SVG, PDF and LATEX format and the scaffold in extended Newick format, and stores complete analyses, including the tree-membership annotations, in a text-based document format. Installers for macOS, Windows and Linux are provided. All figures in this article were produced with version 1.1.2 of the PhyloParallelograms software.

### Statistics and reproducibility

No statistical hypothesis tests were performed. The synthetic comparison (Fig. 3e) is based on 100 independent tree pairs for each of 17 Robinson–Foulds distances, all of which were analysed; box plots show the median and interquartile range, with whiskers extending to the minimum and maximum. Both quantities compared are deterministic functions of a tree pair, so that repeated computation yields identical results.

## Data availability

The tree files used to produce all figures are available at https://github.com/husonlab/phyloparallelograms/tree/main/publication-data. The *Anopheles* alignments, locus coordinates and gene trees are available at https://github.com/husonlab/trees-to-networks-tutorial; the underlying whole-genome alignment is available from Dryad (doi:10.5061/dryad.f4114). The synthetic tree pairs were taken from ref. 35. The species tree and whole-genome tree of Fig. 1 were transcribed from Fontaine et al. ^7^, the cat trees from Li et al. ^11^, the Danthonioideae trees from Pirie et al. ^13^, and the Fagaceae trees were recomputed from data provided by Zhou et al. ^12^ on Dryad (https://doi.org/10.5061/dryad.vq83bk3tc). Source data for Fig. 3e are provided at the URL provided at the beginning this section.

## Code availability

PhyloParallelograms is open-source software released under the GNU General Public License v3. Source code, installers for Linux, macOS and Windows, the user manual and example data sets are available at https://github.com/husonlab/phyloparallelograms. The scripts used for locus selection and extraction from the *Anopheles* alignments are available at https://github.com/husonlab/trees-to-networks-tutorial. The scripts used for the synthetic comparison (batch computation of displacement and hybridization number) are available in https://github.com/husonlab/phyloparallelograms/tree/main/publication-data/figure3.

## Acknowledgements

L.Z. was supported by the Singapore MOE Academic Research Fund Tier 1 [A-8001951-00-00].

D.H.H. and B.C. were supported by institutional funds.

## Author contributions

D.H.H., B.C. and L.Z. developed the concept and contributed to the algorithms. D.H.H. wrote the software and drafted the manuscript. B.C. implemented parts of the algorithm and inferred the Fagaceae trees. D.H.H. prepared the *Anopheles* data and ran the synthetic comparison. All authors edited and approved the manuscript.

## Additional information

**Correspondence** and requests for materials should be addressed to Daniel H. Huson.

